# A post-mitotic *in vitro* murine as a model of muscle damage and repair

**DOI:** 10.1101/2024.10.04.616603

**Authors:** Angelo Galluccio, Samantha Maurotti, Francesca Rita Noto, Francesca Scionti, Carmelo Pujia, Elisa Mazza, Yvelise Ferro, Rosario Mare, Nadia Geirola, Bernadette Scopacasa, Patrizio Candeloro, Luca Tirinato, Angela Sciacqua, Arturo Pujia, Stefano Romeo, Tiziana Montalcini

## Abstract

Sarcopenia is a degenerative condition characterized by the atrophy and functional decline of myofibers, resulting in disability. While the clinical risk factors are known, there is no validated *in vitro* model to understand the molecular mechanisms and identify therapeutics. To tackle this challenge, we generated an *in vitro* post-mitotic muscular system by differentiating mouse myoblast cells, namely C2C12. After 12 days of differentiation, cells were expressing physiological markers of myotubes and became self-contracting. Importantly, transcriptomic analyses demonstrated high similarity (r=0.70) when compared to primary human myotubes (HSkMC) providing evidence of resemblance to human cells. Next, we starved and incubated cells with dexamethasone and observed myotube shrinkage, oxidative stress, modification of anabolic, inflammatory, and catabolic markers recapitulating sarcopenia. Conversely, cell refeeding resulted in a recovery in the model with nutrient deprivation but not when incubated also with dexamethasone. In conclusion, we present a model of sarcopenia due to nutrient deprivation and corticosteroids. This model may allow more efficient and effective future research to identify therapeutics against sarcopenia in humans.

## INTRODUCTION

Sarcopenia is characterized by systematic muscle degeneration, resulting in a progressive reduction in muscle mass, quality, and strength in the skeletal muscle ^1^. Individuals affected by sarcopenia experience a decline in muscle strength, which impairs their mobility and significantly increases the risk of bone fractures from accidental falls, making sarcopenia a major health concern in Western countries ^1^. Indeed, an estimate of the economic burden of sarcopenia is several billion dollars per year in the United States in 2024 ^2^.

Moreover, individuals with sarcopenia have a higher risk of metabolic disorders raising their mortality risk ^3^. Clinical risk factors for sarcopenia are well-defined ^4^. Among these, aging ^5^, malnutrition^6^, and exogenous/endogenous corticosteroid^7,8^ are the most common risk factors for sarcopenia.

Sarcopenia involves various biological mechanisms contributing to the loss of muscle mass in humans^1^. Among these, muscle atrophy leads to a reduction in the number and size of muscle fibers, especially the fast-twitch type II fibers^9,10^. This condition is exacerbated by reduced protein synthesis and increased catabolism, worsening protein balance and resulting in progressive muscle mass loss. Oxidative stress plays a significant role, with the accumulation of reactive oxygen species (ROS) damaging proteins, lipids, and DNA in muscle cells, eventually leading to damage and loss of function^11^.

Corticosteroids contribute to sarcopenia by breaking down the heavy chain of myosin, a key protein involved in muscle contraction^12^. This degradation is accompanied by a reduction in the protein synthesis. This occurs as corticosteroids block the signaling for protein synthesis via the Akt/mTOR pathway^13^. Additionally, a commonly used corticosteroid, dexamethasone, has been found to enhance the ubiquitin-proteasome mediated degradation^14^ by increasing the expression of MAFbx and MuRF1^15,16^.

This chain of events occurs in the muscle tissue, which is composed of multinucleated, post-mitotic muscle cells specialized in generating contractile force^11^. Another fundamental aspect is the role of satellite cells, which are adult muscle stem cells that repair and maintain post-mitotic skeletal muscle tissue. These cells are located between the basal lamina and the sarcolemma of muscle fibers normally in a quiescent state. After injury satellite cells become activated, proliferate, and differentiate into myocytes, which fuse to generate new muscle tissue^17–19^. Studies have shown a strong correlation between the loss and dysfunction of satellite cells and sarcopenia^20–22^.

There is no standardized *in vitro* model of human sarcopenia to understand the molecular mechanisms and identify potential treatments for sarcopenia. In fact, despite the presence of numerous *in vitro* models of skeletal muscle atrophy, these models are not validated and questions remain on their translation in humans^23^. In this context, there is controversy on the duration of myogenic differentiation in C2C12 cells which are commonly used to study muscle atrophy. Differentiation times are defined arbitrarily raising concerns about the validity, reproducibility, and translational of these studies.

Therefore, we sought to develop a model of muscle differentiation by using murine myoblasts that we validated against human primary myocytes. This model was exposed to nutrient deprivation and corticosteroids to induce sarcopenia and subsequently reversed by refeeding.

## MATERIAL AND METHODS

### Cell Cultures and Maintenance

Mouse myoblast C2C12 cell line and human skeletal muscle HSkMC cells were purchased from the American Type Culture Collection (ATCC, Manassas, VA).

C2C12 cells were cultured in 100 mm tissue culture disks (Corning, Kennebunk, USA) and maintained in Dulbecco’s Modified Eagle Medium High Glucose (DMEM) (Corning, Kennebunk, USA) growth medium (GM), supplemented with 10% Fetal Bovine Serum (FBS) (SIAL, Rome, Italy) and 1% penicillin-streptomycin (100μg/ml) (P/S) (SIAL, Rome, Italy) at 37 °C under a 5% CO_2_humidified atmosphere. Cells were sub-cultured when they reached 70 to 80% of confluence. Only cells between passages 3-10 were used in this study. Cells were routinely tested for mycoplasma assay.

HSkMC cells were cultured in 100 mm tissue culture disks (Corning, Kennebunk, USA) and maintained in the Mesenchymal Stem Cell Basal Medium (ATCC, Manassas, VA) and Primary Skeletal Cell Muscle Growth Kit (GM) (ATCC, Manassas, VA) at 37 °C under a humidified atmosphere of 5% CO_2_. Cells were sub-cultured when they reached 80 to 90% of confluence. Cells from passages 1 to 3 were used in this research.

### Differentiation from myoblasts C2C12 to post-mitotic myotubes

C2C12 cells were cultured in p60 (Corning, Kennebunk, USA) at a density of 200.000 cells. After reaching 80–90% confluence (48 hrs), C2C12 cells were rinsed with phosphate-buffered saline (PBS) (Corning, Kennebunk, USA) and the GM was replaced with a differentiation medium (DMEM high glucose) containing 2% horse serum (HS) (Gibco by ThermoFisher Scientific), 1% P/S. This was done to promote the fusion of myoblasts into myotubes over a time period of 0, 3, 6, 9, and 12 days, with the aim of determining the optimal time for mature myotube formation.

### Differentiation from HSkMC cells to myotubes

HSkMC cells were seeded onto a fibronectin-coated p60 at the density of 50.000 cells/cm^2^ and incubated for 24 hours. When cells reached 80–90% confluence, they were rinsed with PBS and the GM was replaced with the DM for 120 hours. Fresh DM was replaced every 48 hours. Primary cell morphology was observed by an Axio Vert.A1 inverted microscope (Carl Zeiss AG).

### Induction of sarcopenia and reverse of the phenotype

To induce sarcopenic damage, the C2C12 myotubes, after 12 days of differentiation, were treated with either phosphate-buffered saline (PBS) for 4 hours or serum-free DMEM containing high glucose and 100 μM dexamethasone (DEX) (Acros organics, New Jersey, USA) for 24-48 hours. To induce the reversal of the damaged phenotype, the culture medium was restored with differentiation medium (DMEM high glucose) containing 2% HS for 120 hours. The differentiating medium was replaced daily.

### Hematoxylin and eosin staining and measurement of myotube diameters

C2C12 cells at 0, 3, 6, 9, and 12 days, following treatment with either PBS for 4 hours or 100 µM DEX for 24 or 48 hours, were first washed twice with PBS and then fixed in 4% PFA for 10 min. Subsequently, the myotubes were stained with hematoxylin and eosin (H&E) solution (BioOptica) for 1 min at room temperature and observed using an optical microscope (Leica, Wetzlar, Germany). For each condition, six images were randomly captured from each well of the six-well plates using a Leica DM4 B Upright Microscope (Leica Microsystems), using an HC FL PLAN 40x/0.65 objective (Corning, Kennebunk, USA). The diameters of three different sites in each myotube were measured using ImageJ software, and at least 120 myotubes in one well were measured.

### Protein extraction and western blot analysis

C2C12 protein lysates, collected at 0, 3, 6, 9, and 12 days, and after treatment with either PBS for 4 hours or 100 µM DEX for 24 and 48 hours and refeeding, and HSkMC protein lysate collected at 0 and 120 hours, were prepared using M-PER protein extraction reagent (ThermoFisher Scientific) supplemented with protease (Roche, Mannheim, Germany) and phosphatase inhibitor cocktails (ThermoFisher Scientific), and briefly homogenized. Samples were loaded into 10% polyacrylamide gels with the PAGE system (BioRad, USA) and run in the SDS running buffer (25 mM Tris, 192 mM glycine, 0.1% SDS, pH 8.8). The run was monitored following ladders: Biorad Precision Plus Protein Standards #1610394 or Thermo Fisher Page Ruler Prestained Protein Ladder #26616. Afterward, proteins were transferred to TransBlot ® Turbo™ Midi-Size Nitrocellulose (BioRad, USA) with Trans-Blot® Turbo™ Transfer System (BioRad, USA) in TransBlot® Turbo™ 5x transfer buffer (BioRad, USA). Membranes were blocked in 5% BSA powder in TBST (25 mM Tris, 150 mM NaCl, 0.2% Tween-20 (Sigma), pH 7.4 adjusted with HCl) and incubated overnight with the indicated primary antibodies dissolved in TBST containing 5% BSA.

Mouse anti-MyHC (Merck Millipore Ltd, Ireland; diluition 1:1000), mouse anti-actinin (Sigma-Aldrich, St. Louis; diluition 1:1000), rabbit anti-pAmpkα (Cell Signalling, United States; diluition 1:1000), rabbit anti-p-Akt (Cell Signalling, United States; diluition 1:1000), rabbit anti-NF-kB (Cell Signalling, United States; diluition 1:1000), rabbit anti-Murf1 (Thermo Fisher Scientific; diluition 1:1000), rabbit anti-Atrogin1 (Abcam, UK), rabbit anti-pErk 1/2 (Cell Signalling, United States; diluition 1:1000), rabbit anti-MyoD1 (Sigma-Aldrich, St. Louis; diluition 1:1000) and mouse anti-β-Actin (Sigma-Aldrich, St. Louis; diluition 1:10000), mouse anti-CALNEXIN (Thermo Fisher Scientific diluition; 1:5000) primary antibodies were used. Blots were probed with primary antibodies, followed by the appropriate horseradish peroxidase (HRP)-conjugated secondary antibody (Bio-Rad, USA), and developed using Clarity™ Western ECL substrate (BioRad, USA). The image captures and densitometric analyses were performed with the ChemiDoc MP Imaging system (BioRad, USA) and ImageJ software, respectively. Uncropped scans of western blots are shown in supplementary information (Source Data_ Unprocessed Western Blot)

### RNA extraction and quantitative reverse transcriptase-PCR

Total RNA from C2C12 cells, at 0, 3, 6, 9, and 12 days, and after treatment with either PBS for 4 hours or 100 µM DEX for 24 and 48 hours, was extracted using TRIzol reagent (Thermo Fisher Scientific) following the manufacturer’s instructions. cDNA was synthesized from 1 µg of total RNA using a high-capacity cDNA reverse transcription kit (Applied Biosystems by Thermo Fisher Scientific). The mRNA expression of *Bmp4, Miogenin, Myf5, Myod-1, Mrf4, MyHCI, MyHCIIa, MyHCIIb, MyHCIIx, Pax-3 and Pax-7* were quantified through Real-Time PCR using SYBR® Green dye (BioRad, USA) (Supplementary Table 1). Data were analyzed using the 2−ΔΔ^Ct^ method and normalized to *β-Actin*.

### RNA-seq analysis

Total RNA from differentiated HSkMC and C2C12 myotubes after 120 hours and 12 days of differentiation, respectively, was extracted with the RNeasy Plus mini kit (Qiagen, Hilden, Germany). RNA integrity number and concentration were evaluated by the Agilent Bioanalyzer 2100 system (Agilent Technologies, CA, USA) and all samples had a RIN>6.8 and concentration >20 ng/μL. Illumina sequencing and quality control analyses were performed by Biomarker Technologies (BMKGene) GmbH (Germany). Reference genome and gene model annotation files for human (GRCh38/hg38) and mouse (GRCm38/mm10) were downloaded from the GENCODE website. Indexes of the reference genome were built using STAR and paired-end clean reads were aligned to the reference genome using STAR aligner (v2.7.11a). Read counting was performed by StringTie (v2.2.1). The Fragments Per Kilobase of transcript per Million fragments mapped (FPKM) were calculated According to the gene length and read counts mapped to a gene. The list of orthologous transcripts in human and mouse (Supplementary Table 2) was extracted from the Ensembl database using the BioMart tool. Further statistical analyses were performed with R software. For pairwise Pearson analysis, a correlation matrix of log-transformed FPKM values was calculated and input into the R function.

### Reactive oxygen species assay

To evaluate reactive oxygen species (ROS), C2C12 myotubes under different conditions were fixed in 4% PFA for 15 min. Subsequently, myotubes were stained using the DCFDA/H2DCFDA Cellular ROS Assay Kit (Abcam, UK) according to the manufacturer’s instructions. ROS generation was observed using a FLoid™ Cell Imaging Station (Thermo Fisher Scientific) at 20× magnification.

### Raman Measurements

C2C12 myoblasts and their differentiated myotube counterparts, following a 12-day differentiation period, were seeded onto CaF_2_ substrates because of their negligible Raman signal, and subsequently cultured in either GM or DM.

Before carrying out Raman analysis, the culture medium was aspirated, and cells were washed three times with pre-warmed PBS. This was followed by a fixation process using 4% formalin for 15 min at room temperature. Soon after, cells were rinsed with PBS to eliminate residual fixative.

Raman micro-spectroscopy was performed using a Witec Alpha-300RA system equipped with a 532 nm wavelength laser source. The laser, operating at 10 mW/cm², was focused on the sample through a 50X/0.75 NA objective in air. The laser beam was automatically scanned over fixed cells in a grid pattern with 0.4 µm steps, which is comparable with the optical diffraction limit of the setup. An integration time of 1 sec per spectrum was used, thereby assembling a hyperspectral dataset with one spectrum obtained for each pixel in the scanned area of the cellular sample.

### Raman Spectra Processing

Before analysis, the acquired Raman datasets underwent preprocessing to ensure comparability across measurements taken at different times on various cell cultures. Initially, baseline correction was applied to each single-cell dataset, involving the subtraction of a first-order linear fit within the high-frequency region (2600-3100 cm⁻¹) and a fourth-order polynomial in the low-frequency, fingerprint region (400-1800 cm⁻¹). Following this, normalization was executed on each Raman map, using the maximum spectral area recorded on the map as normalization factor, thus ensuring standardized signal intensities across different maps. For collective spectral feature assessment within the same cell treatment groups, principal component analysis (PCA) and K-means clustering analysis (KCA) were employed. The preprocessing and subsequent analytical procedures were conducted using the freely available Raman Tool Set software package ^24^.

### Statistical Analysis

The data are presented as mean ± standard deviation (SD) from at least three independent experiments and were analyzed using two-tailed Student’s t-test and linear regression. Values with p<0.05 were considered statistically significant. Statistical analysis was performed using GraphPad Prism 9.3.1.

## RESULTS

### Myoblasts differentiate into myotubes after 12 days

The study design is illustrated in Figure 1. To generate an *in vitro* post-mitotic muscle model, mouse C2C12 cells were differentiated over a time course of 0, 3, 6, 9, and 12 days (Figure 2). First, we measured *Pax3* and *Pax7* gene expression as markers of initiation of myogenic differentiation. *Pax3* expression was higher after 3 days as compared to day 0 and became lower in a time-dependent manner after 12 days of differentiation (Figure 2b) (PMID: 7744814). *Pax7* levels were similar to *Pax3* except for day 12 where *Pax7* levels were higher. (Figure 2c). The increase in *Pax7* on day 12 suggests the presence of quiescent progenitor cells^25,26^.

**Figure 1.**
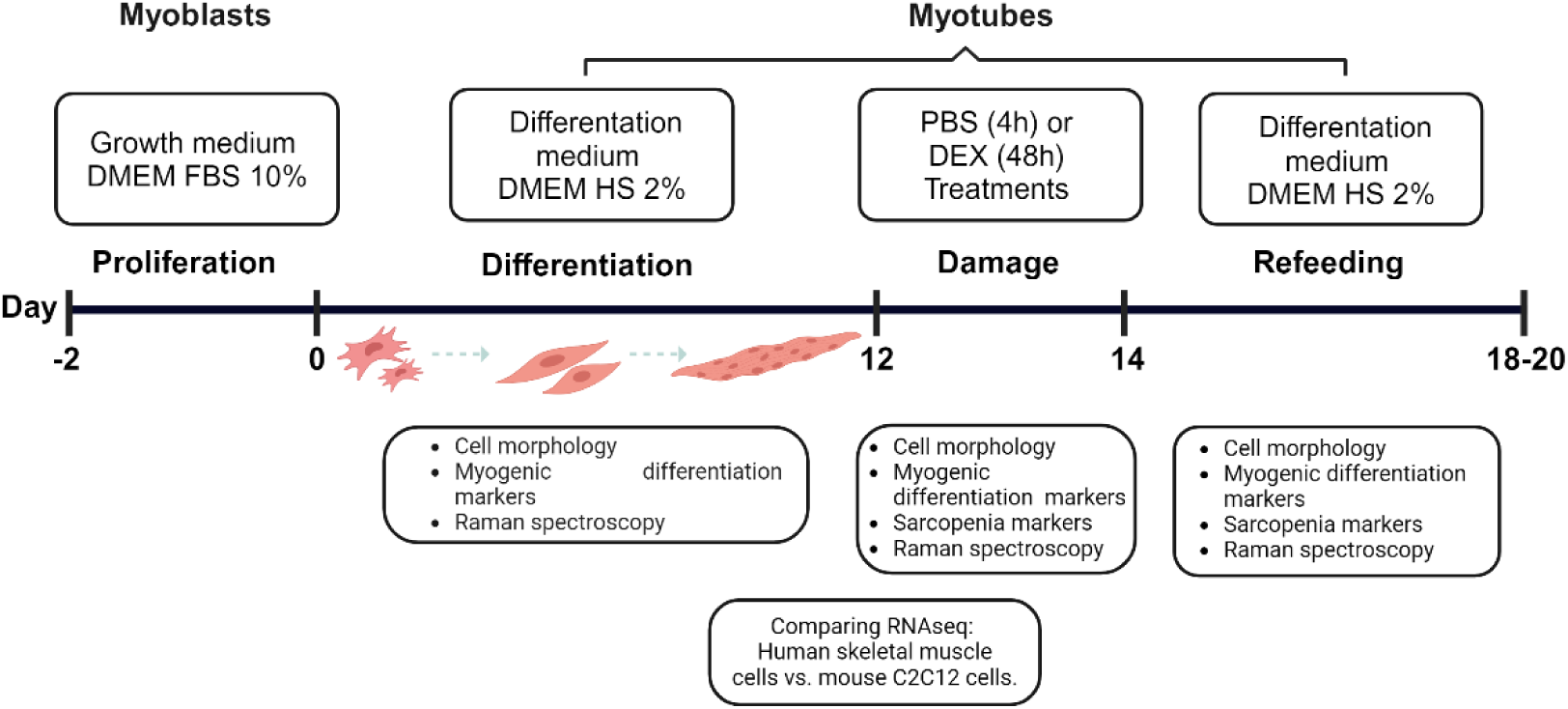
Experimental design of the study.

**Figure 2.**
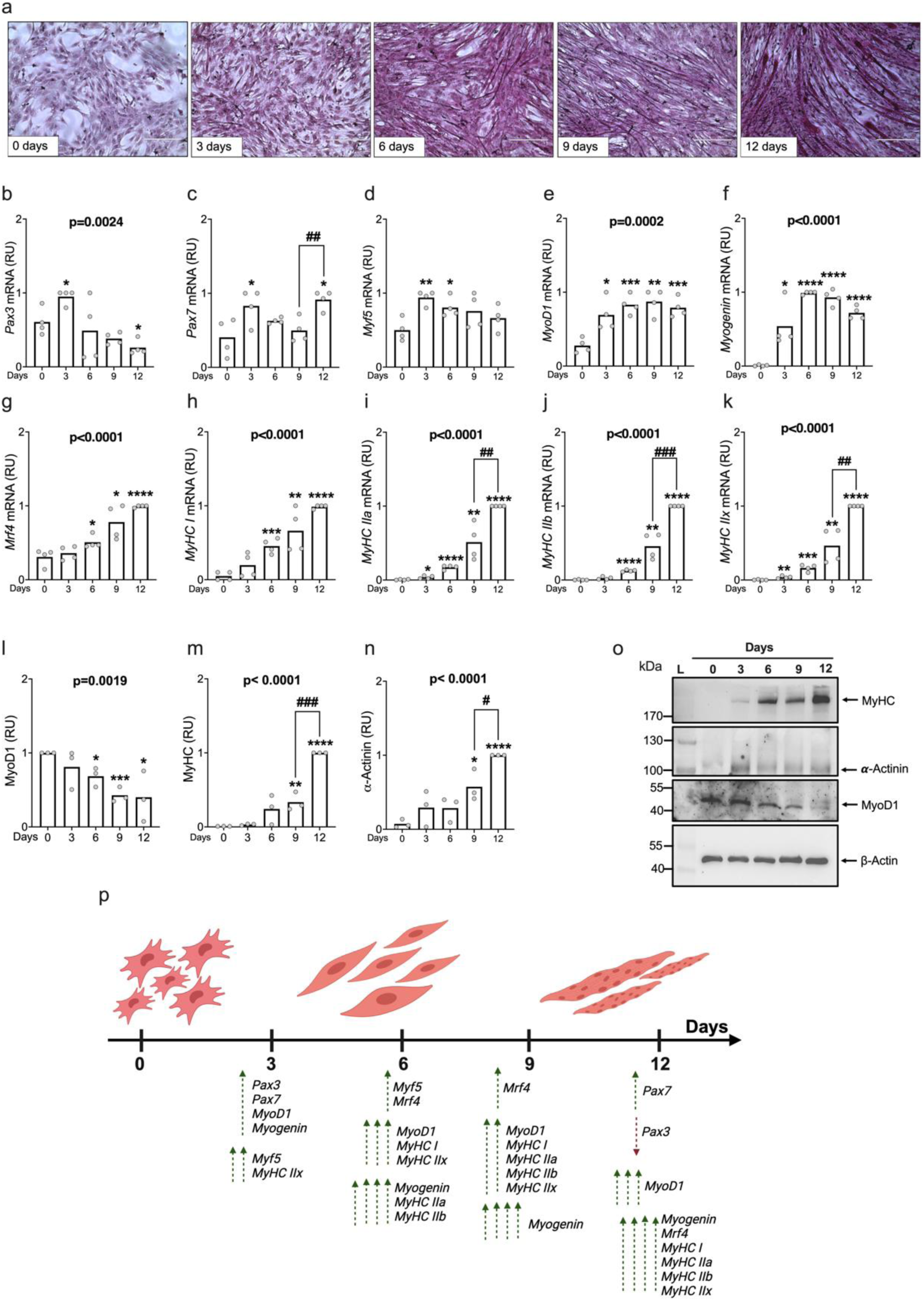
C2C12 myoblast differentiation to post-mitotic myotube over 12 days. **(a**) Representative images of H&E staining (×20 magnification) at day 0, 3, 6, 9, 12; **(b-k)** mRNA expression levels of *Pax3*, *Pax7*, *Myf5*, *MyoD*-1, *Myogenin*, *Mrf4*, *MyHC I*, *MyHc IIa*, *MyHC IIb* and *MyHC IIx* were measured using Real-Time PCR. Data were analyzed using the 2^-ΔΔCt^ method and normalized to β-Actin. **(l-o)** Protein levels of MyHC and α-Actinin, markers of later differentiation to myotube and MyoD-1 protein levels as a marker of early differentiation of the myotubes. **(p)** Quantification of protein levels relative to β-Actin. Protein levels were measured by western blotting (see methods for antibodies). **(p)** Illustration depicting the modulation of genes and proteins involved in various stages of myogenic differentiation. Data are represented as mean ± SD of three independent experiments. *p*-values calculated by Student’s t-test: vs day 0 \**p* < 0.05, \*\**p* < 0.01; ***p<0.001; ****p<0.0001; vs day 9 # *p* < 0.05, ## *p* < 0.01; ### p<0.001; p-values in the figure are calculated by using the oneway anova test for linear trend. Abbreviations: RU= Relative unit.

Next, we measured mRNA levels of genes involved in myogenic differentiation, namely *Myogenic factor 5* (*Myf5*), *Myoblast determining protein 1* (*MyoD1*), *Myogenin* (*Myog*), and *Muscle-Specific Regulatory Factor 4* (*Mrf4*) (Figure 2 d-g). *Myf5* increased on day 3 and continued with a time-dependent decrease afterward, *MyoD1* increased on day 3 and reached a plateau until day 12. These data are consistent with the beginning of myoblast differentiation but not yet fusion. *Myog* increased on day 3 with a plateau on day 6 and then decreased on day 12, indicating myocyte fusion. Finally, *Mrf4* increased from day 6 to day 12 in a time-dependent manner suggesting differentiation of myoblast into myotubes.

Next, we measured markers of late differentiation *Myosin Heavy Chain 1 (MyHC I)*, *Myosin Heavy Chain IIA (MyHC IIa)*, *Myosin Heavy Chain IIb* (*MyHC IIb)*, and *Myosin Heavy Chain IIx (MyHC IIx)* (Figure 2 h-k). All these markers were expressed at the highest level at day 12 consistent with the observation of myotubes contraction at day 12 (Supplementary Video 1). Finally, we examined protein levels of one marker of initial differentiation (MyoD1) and two of late differentiation, namely MyHC and α-Actinin (Figure 2 l-n). Consistently with the mRNA data, MyoD1 had its highest expression at day 3 and then diminished in a time-dependent manner (Figure 2 l). MyHC and α-Actinin were detectable at day 3 and then increased in a time-dependent manner.

Interestingly, there was a discordance between the mRNA and protein levels of MyoD1. Indeed, while *MyoD1* mRNA increased over time, protein levels decreased. This is consistent with the proteasome-mediated degradation of *MyoD1*^27^. Figure 2p represents a schematic of the genes involved in myogenic differentiation for a 12-day time course.

### Transcriptomic analyses reveal similarities between differentiated and primary myotubes

To understand similarities between our model and human primary myotubes, we differentiated human primary myoblasts into myotubes. As expected, after 5 days of differentiation we observed an increase in both α-ACTININ and MYHC, along with a reduction in MYOD1 (Figure 3 b-d). Next, we performed RNA sequencing of these human primary cells and of the murine myotubes. We identified a total of 63,187 human and 48,435 mouse transcripts. After removing unmatched, duplicated, and unexpressed transcripts, 8.365 orthologous transcripts remained (Figure 3f). Pearson pairwise analyses revealed a strong correlation between mRNA levels of these orthologs (r=0.70; p<0.001; Supplementary Table 2 and Figure 3g), indicating that our model shares a similar transcriptomic signature with human primary myotubes.

**Figure 3.**
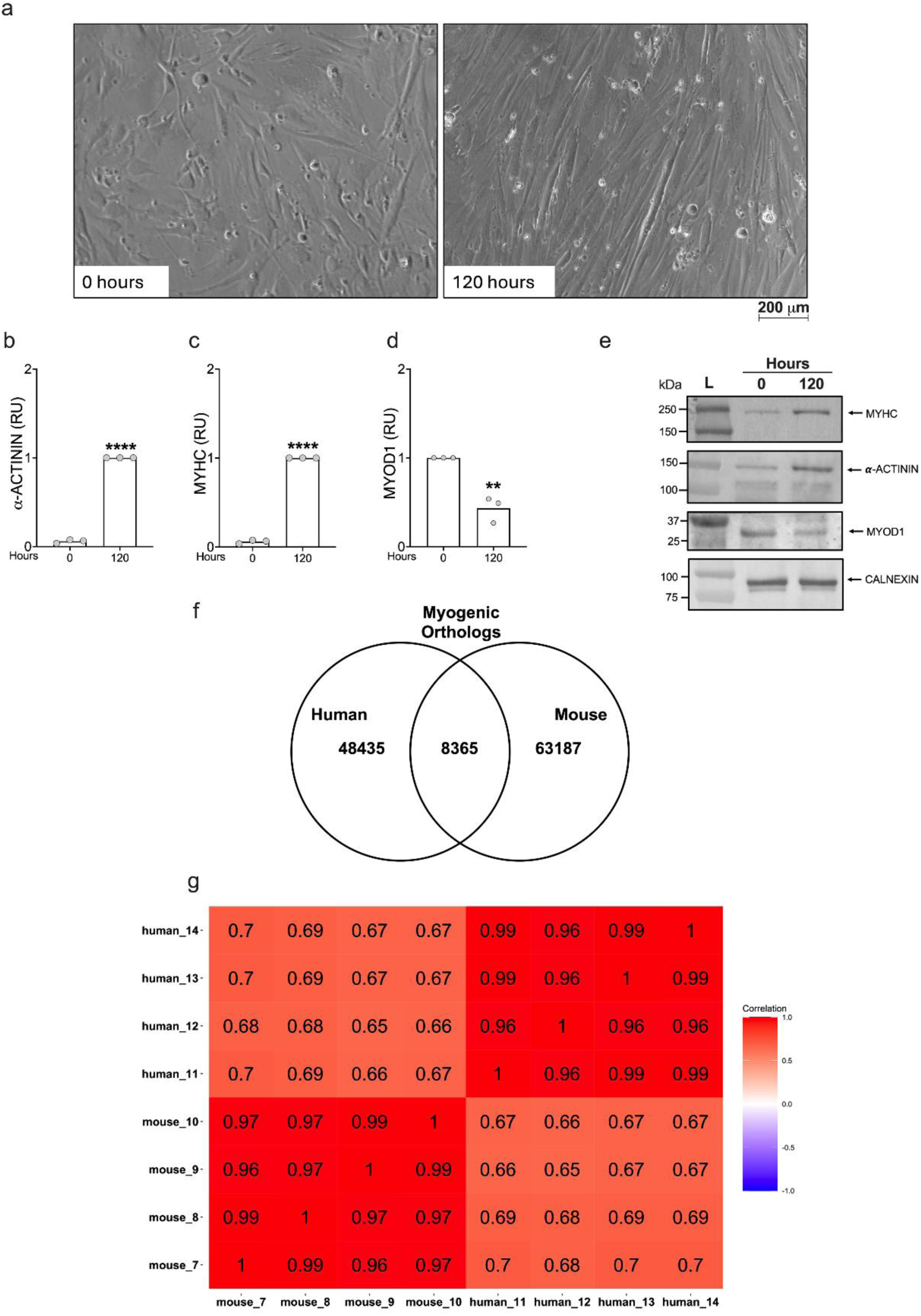
A strong correlation between primary human and mouse differentiated cells. **(a)** Representative images of human HSkMC cells (×20 magnification) by bright field phase-contrast microscope. **(b-d)** MyHC and α-ACTININ and MyOD-1 protein levels in HSkMC cells. **(e)** Quantification of protein levels relative to β-Actin. Protein levels were measured by western blotting (see methods for antibodies). Data are represented as mean ± SD of three independent experiments. p-values calculated by Student’s t-test: **p < 0.01; ****p<0.0001. Abbreviations: RU= Relative unit. **(f)** Transcriptomic analysis revealed a total of 63187 transcripts in humans and 48435 transcripts in mice. After excluding mismatched, duplicated, and unexpressed transcripts, 8365 orthologous transcripts were identified. **(g)** The heatmap demonstrates a robust correlation in the expression patterns of orthologous transcripts between human and mouse differentiation model.

### Induction and reversal of sarcopenia in murine myotubes by nutrient deprivation

Sarcopenia in humans may be due to negative energy balance. Indeed, it is a common clinical observation that negative energy balance reduces adiposity but also muscle mass. Therefore, as a first step, we decided to starve cells by incubating them with PBS for 4 hours. Sarcopenia is associated with lower muscle fiber size, inflammation, mitochondrial dysfunction, and oxidative damage from free radicals^28^.

To mimic sarcopenia due to malnutrition and test if our model is reversible, we incubated murine myotubes with PBS for 4 h and then replaced PBS with the differentiation media for 5 days. In line with the presence of sarcopenia, after 4 hours with PBS we observed a striking reduction in diameter of myotubes that were not able to contract (p=0.001; Figure 4 a, b), a striking 10-fold increase in intracellular reactive oxygen species (ROS) (p=0.03; Supplementary Figure 2a,b), a 3-fold increase in inflammation markers (NF-κB: p<0.01; Figure 4c), a robust and consistent increase in three markers of protein degradation (*p*AMPKα: p<0.01; Murf-1: p<0.01; Atrogin-1: p<0.01; Figure 4 d-f), and a reduction in two anabolic markers (pAkt: p<0.05; pErk1/2: p<0.0; Figure 4 g-h).

**Figure 4.**
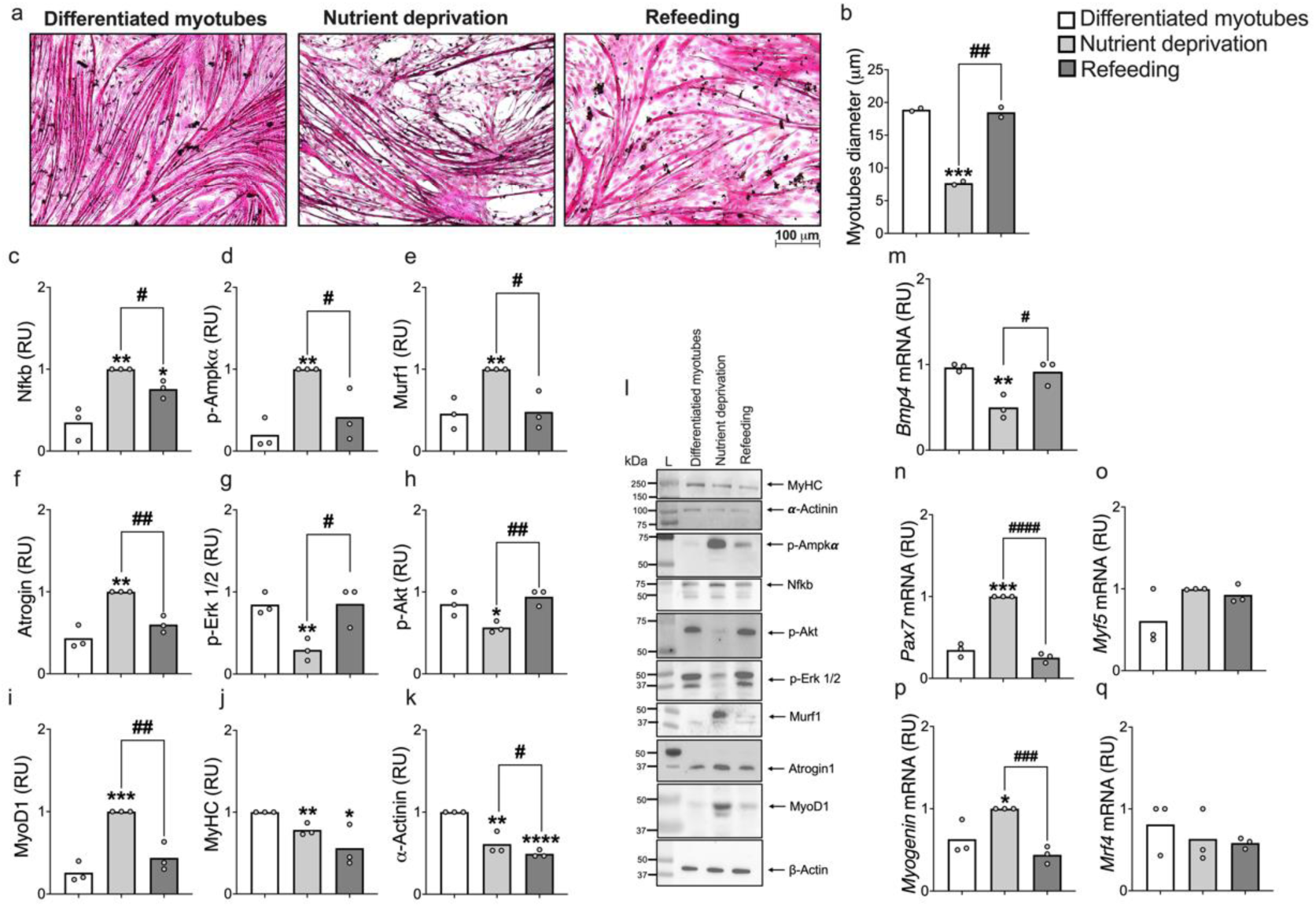
Nutrient Deprivation-Induced Post-Mitotic myotube damage through the increase of catabolic markers, the reduction of anabolic markers with consequent activation of satellite cells. **a-b)** post-mitotic myotubes underwent a 4-hour incubation with phosphate-buffered solution (PBS) before H&E staining (20x magnification), resulting in reduced myotube diameter. Protein levels by western blotting and relative quantification of inflammation **(c)** catabolic/atrophy (**d-f**), anabolic **(g,h)** and differentiation **(i-k)** markers. However, refeeding with differentiation media after 120 hours reversed phenotype, inflammation, catabolic, anabolic and differentiation markers. **(m-q)** mRNA expression levels of *Pax7*, *Myf5*, *MyoD*-1, *Myogenin*, *Mrf4*, were measured using Real-Time PCR. Data were analyzed using the 2^-ΔΔCt^ method and normalized to β-Actin. Data are represented as mean ± SD of three independent experiments and p values are calculated by Student’s t-test: vs Differentiated myotubes \**p* < 0.05, \*\**p* < 0.01; ***p<0.001; ****p<0.0001; vs Nutrient deprivation group # *p* < 0.05, ## *p* < 0.01. Abbreviations: RU= Relative Unit.

After 5 days of refeeding, cells appeared healthier and myotubes diameter returned to baseline together with inflammatory (NF-κB; Figure 4c), protein degradation (*p*AMPKα, Murf-1, and Atrogin-1; Figure 4 d-f), and early differentiation markers (MyoD1; Figure 4i). However, myotubes were still not able to contract and, consistently, we found markers of late differentiation (α-Actinin and MyHC; Figure 4 j, k) that remained reduced.

Regarding early myogenic markers, we observed that the levels of *Pax7* and *Myogenin* are increased post-injury (Figures 4 n,p); while no changes were observed in the expression levels of *Myf5* and *Mrf4* (Figures 4 o,q). Conversely, *Bmp4*, a marker of quiescence of satellite cells, was reduced by the injury. After refeeding, there was a reduction in *Pax7* and *Myogenin (Figures 4 n, p)*, and an increase in *Bmp4 (Figure 4 m)* that is consistent with the initiation of the repair mediated by the activation of satellite cells.

### Induction and reversal of sarcopenia in murine myotubes by excess of dexamethasone and starvation

To mimic sarcopenia due to an excess of glucocorticoids and starvation, we exposed murine myotubes with high doses of dexamethasone (DEX) (100 µM) with a time course (Supplementary Figure 2). Dexamethasone incubation and starvation led to a 50% reduction in myotube diameters (Figure 5a, b) and a time-dependent increase in ROS levels with a 12-fold increase after 48 hours (Supplementary Figure 2 a,b). Additionally, dexamethasone incubation and starvation increased catabolic markers (Murf-1 and Atrogin-1; Figure 5 d,e), reduced anabolic markers (pAkt and pErk1/2, Figure 5g-h) and reduced NF-kB (Figure 5c), a marker of inflammation.

**Figure 5.**
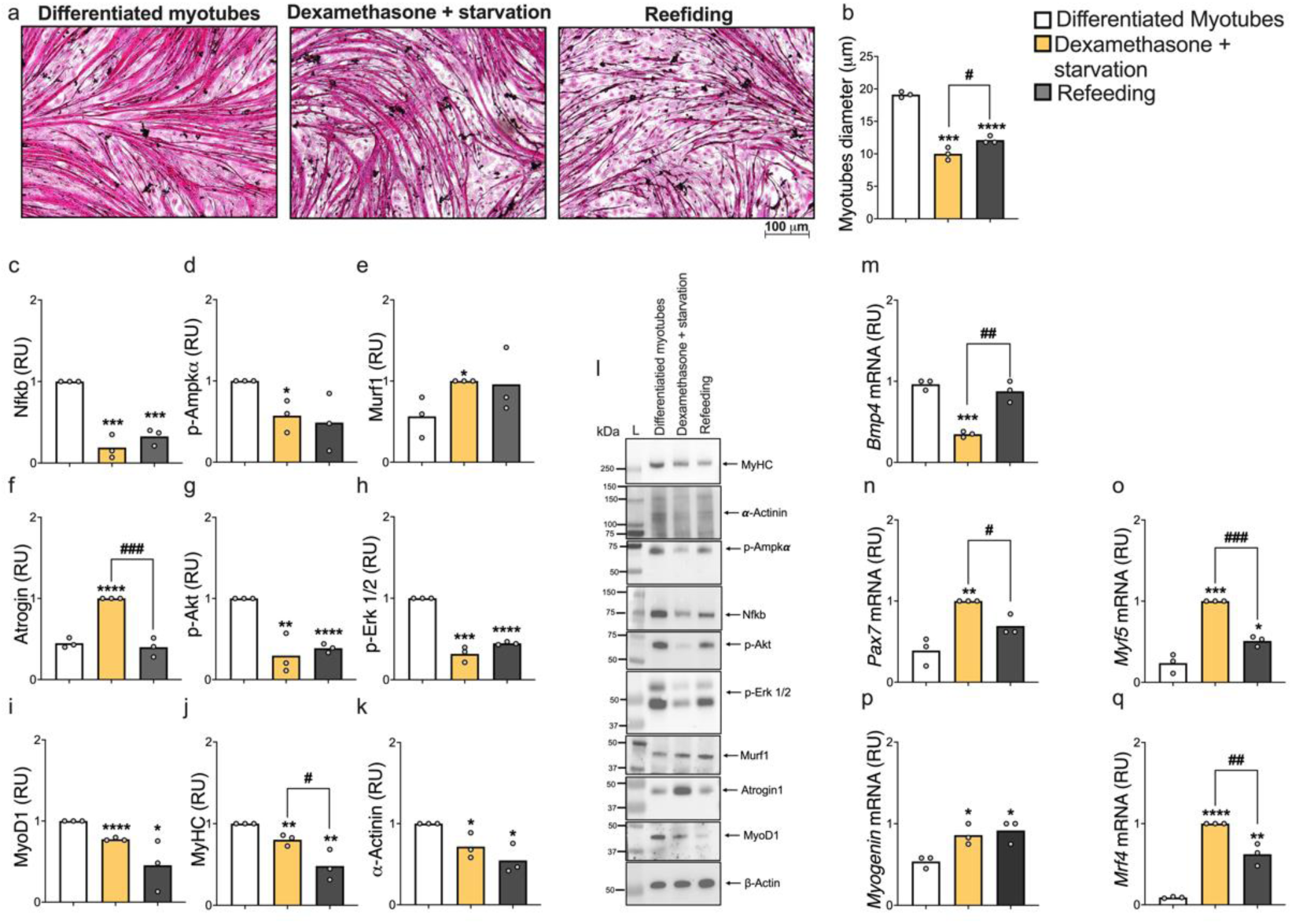
Glucocorticoid-Induced Post-Mitotic Myotube Damage through the modulation of catabolic, anabolic, and differentiation myogenic markers. **a)** Myotubes differentiated for 12 day and incubated for 48 hours with 100 µM dexamethasone in H&E staining (x 20 magnification) **(b)** reduces diameter of myotubes. Protein levels by western blotting and relative quantification of anabolic **(g-h)**, catabolic/atrophy **(d-f)** and differentiation **(i-k)** markers. However, refeeding with differentiation media after 120 hours reverted Atrogin-1 *Pax*-7, *Myf5* and *Myf4* expression. **(m-q)** mRNA expression levels of *Pax7*, *Myf5*, *MyoD*-1, *Myogenin*, *Mrf4*, *MyHC I*, *MyHc IIa*, *MyHC IIb* and *MyHC IIx* were measured using Real-Time PCR. Data were analyzed using the 2^-ΔΔCt^ method and normalized to β-Actin. Data are represented as mean ± SD of three independent experiments and p-values are calculated by Student’s t-test: vs Differentiated myotubes \**p* < 0.05, \*\**p* < 0.01; ***p<0.001; ****p<0.0001; vs Nutrient deprivation group # *p* < 0.05, ## *p* < 0.01; ### p<0.001. Abbreviations: RU= Relative Unit.

Moreover, markers of early and late contraction (MyoD1, MyHC, and α-Actinin) were reduced (Figure 5 i-k), while gene levels of various *MyHC* isoforms were increased. Regarding the quiescence cell marker *Pax-7*, dexamethasone incubation increased its expression levels (Figure 5n) mirrored by a decrease in *Bmp4* (Figure 5n) suggesting activation of satellite cells. For myogenic differentiation markers, *Myf5*, *Myogenin*, and *Mrf4* were increased (Figure 5 o-q). After 5 days of refeeding, there was an incomplete reversal of the cellular phenotype after incubation with Dexamethasone, and the cell diameter remained lower as compared to baseline (Figure 5a, 5b, and Supplementary Figure 3k). However, the levels of Atrogin-1, *Pax*-7, *Myf5*, and *Mrf4* returned to baseline (Figures 5 f-q). Interestingly, MyHC, MyoD1, and α-Actinin became lower than baseline (Figures 5 i-k).

### Raman spectroscopy shows higher myotubes oxidative phosphorylation that is reduced after damage

To gain a deeper understanding of the intracellular composition and oxidative phosphorylation, we performed Raman spectroscopy on myoblasts, myotubes, and myotubes subjected to nutrient deprivation, dexamethasone treatment, and refeeding (Figure 6). The Raman spectra were different for each of the conditions we examined (Figure 6 a-e, f-j and Source Data_Raman Spectroscopy).

**Figure 6:**
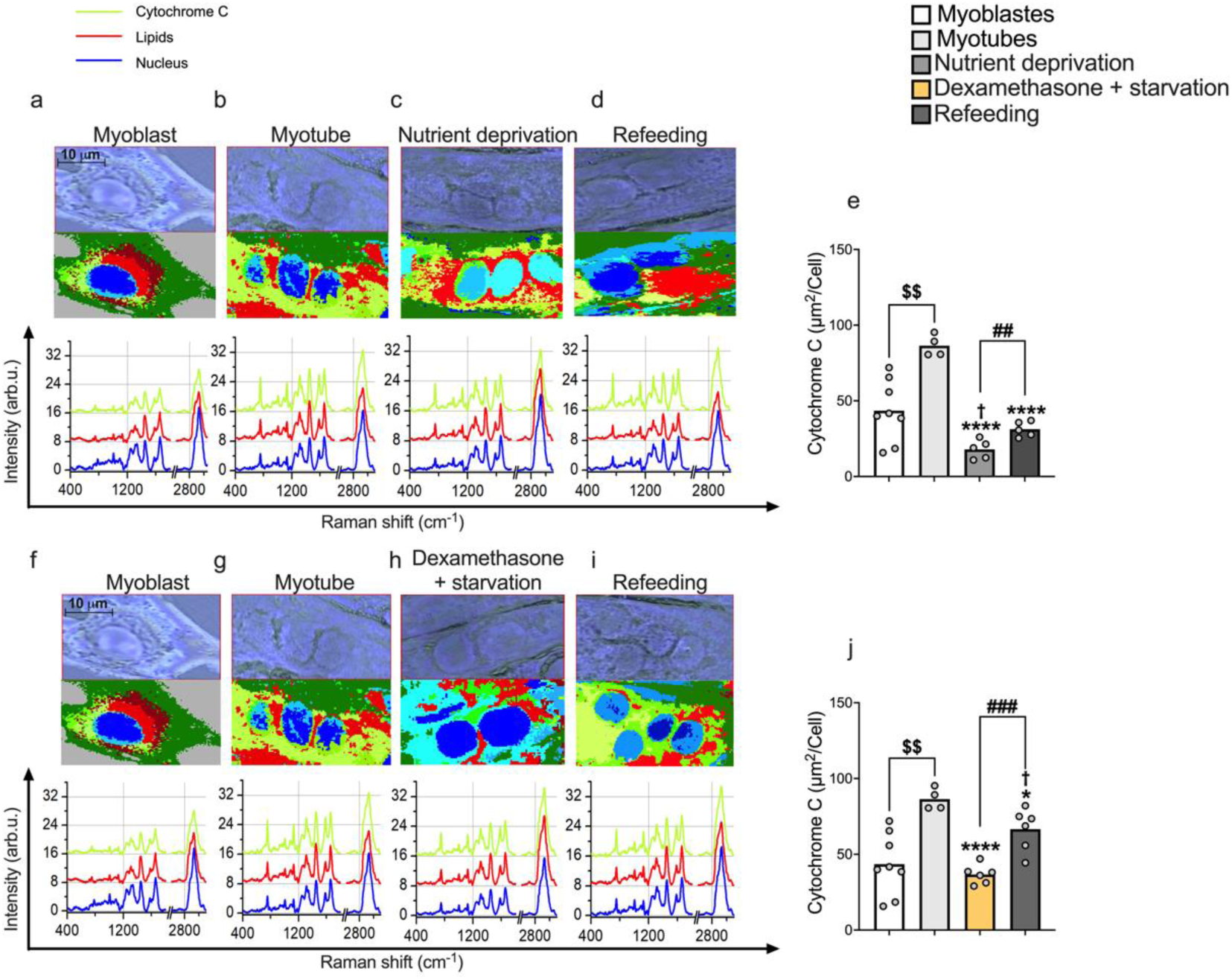
Raman Multivariate Analysis Results. **(a-i)** Showing the comparison between the optical image of the cell acquired in brightfield with the respective map obtained from the KCA. Each map highlights the various cell components such as nucleus (blue, light-blue, cyan), cytochrome C (lime, yellow-green, light-green) and membrane (olive). Graphs above show the average spectra of the main and more significative three classes only of cellular components. (e-j) graphic representation of the statistics calculated considering the spectra relating to the cytochrome C. Data are represented as mean ± SD of eight samples measured per treatment and p-values are calculated by Student’s t-test: vs Myotube group *p < 0.05, **** p<0.0001; vs Myoblaste group $$< 0.01; vs Nutrient deprivation group ## p < 0.01; vs Myoblaste group † p < 0.05.

Cytocrome-C (Cyt-C) was identified by 4 peaks at wavenumbers 750, 1126, 1310, and 1585 cm^-1^ (PMID: 23204423), and Cyt-C levels are calculated as the total area assigned to Cyt-C per cell (Cyt-C, Fig. 6 e,j). Cyt-C is required to generate energy during muscle contraction. Consistently with the observation of cell contraction, Cyt-C increased after myotube differentiation. Damage induced by nutrient deprivation and dexamethasone resulted in a reduction in Cyt-C as compared to the myotube and the myoblast. Moreover, refeeding increased the Cyt-C levels as compared to the myoblast but remained lower as compared to the myotubes. This is consistent with the absence of contraction after refeeding in both models.

## Discussion

In this work was established a protocol for differentiating murine myoblast into myotubes, we validated it against human primary myocytes, and we subsequently used it to induce sarcopenia by starvation and corticosteroid incubation.

We started by differentiating murine myoblast into myotubes after 12 days. These myotubes display all the salient features of postmitotic muscle tissue including high levels of all myosin isoforms. Moreover, we found high levels of cytochrome C, a key marker of energy production eventually leading to cell contraction^29^. Consistently, the differentiation culminated with the ability of myotubes to contract. Moreover, we observed the presence of satellite cells which are key cells after injury for muscle regeneration^30^.

Specifically, we have demonstrated that after a differentiation period of 12 days, there is an increase in Pax7 expression, suggesting a quiescent satellite cell state^31^. Furthermore, we observed that following muscle damage, the rise in this marker, coupled with a decrease in *Bmp4* expression, indicates the activation of satellite cells^31^.

To validate our murine model in humans, we performed RNAseq and examined the similarities between our model and primary human myocytes. Transcriptomic analyses showed a high correlation among mRNA levels between the mouse and human primary myotubes indicating that our murine model recapitulates the human phenotype.

Sarcopenia is often due to nutrient deprivation and elevated corticosteroid levels^32^. To mimic sarcopenia due to these conditions, we deprived differentiated myotubes of nutrients alone and with incubation of dexamethasone resulting in a) the absence of contraction, b) a consistent reduction in myosin, actinin, cytochrome C, c) reduction in myotube diameter, d) a consistent increase in atrophy and, e) a reduction in anabolic markers.

While with the nutrient deprivation, there was an increase in inflammatory markers^33^, as expected these markers were reduced when myotubes were incubated also with dexamethasone. This finding is supported by Matsumura et al., who demonstrated that dexamethasone inhibits the activation of NF-kB in vascular smooth muscle cells^34^.

Incubation with dexamethasone associated with starvation resulted in more severe and permanent damage as shown by the reduction in MyoD1 that was paralleled by the absence of myoblasts as opposed to an increase in MyoD1 and the presence of several myoblasts during starvation only. This is consistent with the observation in humans that nutrient deprivation associated with elevated levels of corticosteroids results in a more aggressive form of sarcopenia^35,36^. Taken all this together, our data demonstrate that our *in vitro* model mimics sarcopenia. Therefore, one may envisage using this model as a first step to identify therapeutics to prevent sarcopenia before going into primary human cells and *in vivo*.

When we attempted to reverse the sarcopenia phenotype by refeeding the model, myotube diameter, atrophy, anabolic and catabolic markers returned to baseline after nutrient deprivation^32^ but not after incubation with dexamethasone, except for Atrogin-1 that returned to baseline. Therefore, the model of nutrient deprivation and dexamethasone may be also used to identify treatments against sarcopenia.

However, in both conditions, there was no contractility indicating the presence of residual damage also in the refeeding after nutrient deprivation. One may speculate that a longer time of recovery would have allowed a complete restoration of the model.

A strength of our study is that our model recapitulates the human phenotypes in a more affordable model that may be used to study sarcopenia and identify therapeutics.

In conclusion, we present a model of sarcopenia due to nutrient deprivation and corticosteroids. This model may allow more efficient and effective future research to identify therapeutics against sarcopenia in humans.

## Funding

No founding

## Competing interests

S.R. has been consulting for AstraZeneca, GSK, Celgene Corporation, Ribo-cure AB and Pfizer in the last 5 years and received the research grant from AstraZeneca. The funders had no role in the design of the study; in the collection, analyses, or interpretation of data; in the writing of the manuscript, or in the decision to publish the results.

## Figure legend

**Supplementary Figure 1. Nutrient Deprivation-Induced Post-Mitotic myotube damage through the increase of ROS accumulation. (a-b)** post-mitotic myotubes underwent a 4-hour incubation with phosphate-buffered solution (PBS) before ROS staining (20x magnification), resulting in ROS accumulation in nutrient-deprived myotubes. **c**) optimal timing for refeeding after a period of fasting.

**Supplementary Figure 2. Effect of 24- and 48-Hours Glucocorticoid Exposure on Catabolic and Anabolic Markers and ROS Accumulation in Post-Mitotic Myotubes. (a-b)** Myotubes differentiated for 12 day and incubated for 24 or 48 hours with 100 uM dexamethasone before ROS staining (x 20 magnification), resulting in ROS accumulation (n=3). Protein levels by western blotting and relative quantification of anabolic **“c-d”,** catabolic/atrophy **“e-f”,** differentiation **“g”** and inflammation **“h”** markers (n=3). **i)** Quantification of protein levels relative to β-Actin. Protein levels were measured by western blotting (see methods for antibodies). Data are represented as mean ± SD of three independent experiments. *p*-values calculated by Student’s t-test: vs 0 hours * *p* < 0.05, ** *p* < 0.01; ***p<0.001; vs 24 hours * *p* < 0.05 ; p-values in the figure are calculated by using the oneway ANOVA test for linear trend. Abbreviations: NF-kB= Nuclear factor kappa-light-chain-enhancer of activated B cells; MyHC= Myosin heavy chain; p-Akt= phospho-Protein kinase B; p-Erk= phospho-Extracellular signal-regulated kinases; DEX= Dexamethasone; L=Ladder.

